# Retro-orbital blood sampling elicits short-term reductions in feeding behavior in western fence lizards (*Sceloporus occidentalis*)

**DOI:** 10.64898/2026.07.28.741151

**Authors:** May S. Jiang, H. Bradley Shaffer, David R. Daversa

## Abstract

Retro-orbital blood sampling, a method where the retro-orbital (post-orbital) sinus is punctured to draw blood, is often used in herpetological research given its ease, speed, and presumed minimal negative effect on the subject. Although literature establishing the method claims little to no impact on subjects, there exists little data explicitly testing the behavioral or welfare-related effects of this technique. We conducted a field experiment with western fence lizards (*Sceloporus occidentalis*) to address the potential impacts of retro-orbital blood sampling. Whereas handling the lizards had no demonstrable impact on lizard prey capture and feeding, there were significant short-term increases in time to initiate prey attack (attack latency) and time to successfully capture prey (feeding latency) in *S. occidentalis* immediately following blood draws. These increases diminished within 24 hours. Our experiments provide new evidence that this common blood sampling method reduces lizard feeding efficiency in the short-term. Researchers and welfare advocates may consider incorporating these findings into their research designs to optimize animal welfare and the generation of reliable ecological and behavioral data.

## Introduction

Blood sampling is, ideally, a minimally invasive technique that allows the studies of organismal function at physiological and molecular levels, advancing research in disciplines including environmental endocrinology, immunology, molecular ecology, and epigenetics, among many others. Insights from studies involving blood sampling support the greater understanding of individual responses to environmental conditions, as well as those shaping broader patterns in ecology, behavior, and evolution [1–6]. In studies of reptiles, existing methods of blood sampling include cardiac puncture, decapitation, venipuncture, and retro-orbital blood draw. Unlike other methods, retro-orbital blood draw, where a heparinized microcapillary tube is used to puncture the retro-orbital sinus, allows for repeated blood collection from an individual in the same day, making it particularly advantageous for studying short-term physiological changes in reptiles [2,5,7–9]. Additionally, retro-orbital blood sampling requires no anesthesia and can be used at low temperatures, making it the technique of choice for field studies and animal welfare agencies [10,11].

Despite the wide use of retro-orbital blood sampling, little is known about its impacts [12]. Existing studies on reptiles have yielded conflicting results on the technique’s physiological impacts, with some studies demonstrating elevated hormone levels correlated with stress (corticosterone) while others show no effect [9,12–14]. Yet to date, no formal evaluation of the behavioral or welfare impacts of retro-orbital blood collection has been conducted on this procedure for any reptile.

The aim of this study was to determine whether retro-orbital blood sampling had short term effects on lizard feeding behavior, which we consider to be a reasonable proxy for welfare impacts [15,16]. We studied these effects in the western fence lizard (*Sceloporus occidentalis*), a species with a broad geographical and ecological distribution in western North America ([17,18]; Fig 1). *S. occidentalis* is a key model organism in reptilian ecology, and findings for this species often provide insights that are broadly applicable to other squamate reptiles [6,19]. If retro-orbital blood collection negatively impacts western fence lizard welfare, we should observe increased latency and reduced feeding success following a blood draw.

**Fig 1.**
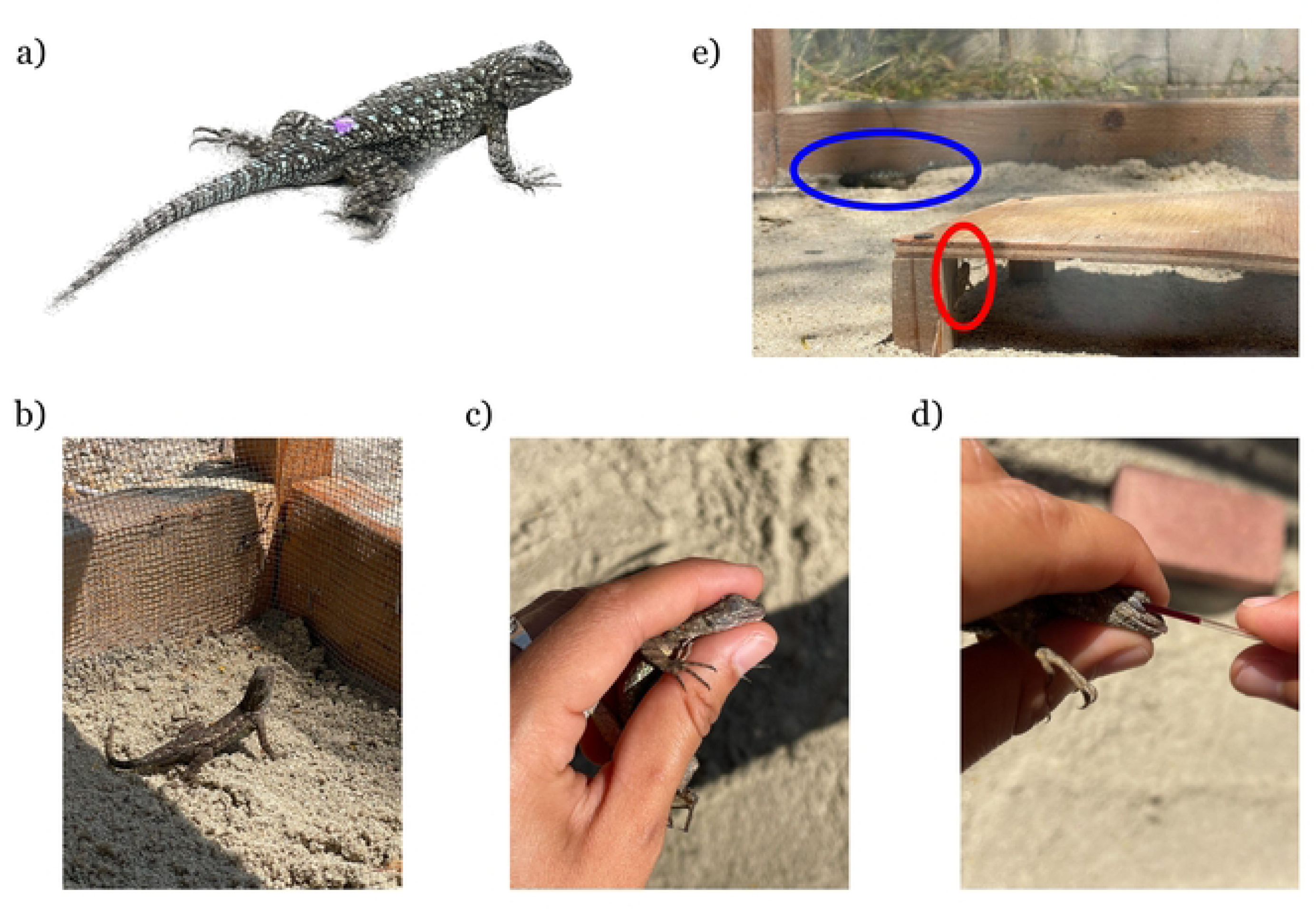
Photo of (a) western fence lizard (*Sceloporus occidentalis*) released with a purple mark to prevent recapture of the same individuals across experiments; (b) An example of an undetectable cricket (circled in red), with the lizard in the background flattened against the sand (circled in blue); (c) control treatment; (d) immobilization; and (e) retro-orbital blood sampling.

## Materials and methods

All procedures were approved by University of California Los Angeles (UCLA) Institutional Animal Care and Use Committee (Protocol No. ARC-2025-016; held by David Daversa and Brad Shaffer). Scientific collecting permits California Department of Fish and Wildlife held by Brad Shaffer (SC 2480). The study period was from May to July 2025.

### Collection and experimental sites

We lassoed eighteen male adult western fence lizards (3 captured per experimental batch, 6 batches total) from sites on the UCLA campus that we previously confirmed had high western fence lizard densities (Jiang 2024, unpublished data). Specific locations within the UCLA campus included: Sunset Canyon Recreation Center (recreational area with irrigated lawns, trees, and garden; latitude/longitude N 34.07531, W -118.4531), Sage Hill (a native chaparral California habitat reserve on the UCLA campus; N 34.07472222, W -118.45611111), and the UCLA Chancellor’s Residence (residential irrigated landscape; N 34.07654, W -118.44215). Upon capture, we recorded ventral body temperature with a laser infrared thermometer (ennoLogic, model et650D), body mass with a spring scale (± 0.5 g; Pesola 20030 Micro-Line Spring Scale 30g x 0.25g), and snout-vent-length (SVL) length from the snout to the mid-cloacal opening with calipers. We used a 50 mm SVL cut off to distinguish adults from juveniles [20], and the presence of enlarged postanal scales to identify males [18]. All lizards were tick-free based on visual inspection. Lizards were placed in clean muslin bags and transported to the experimental site in an empty cooler within 3 hours of capture.

We conducted the experiments at the UCLA Mathias Botanical Garden (N 34.066056, W -118.440322), located roughly 1 km from our capture sites. We held lizards individually in outdoor enclosures (152 cm × 91 cm × 61 cm; length × width × height) built with a latched hinged lid, solid wood base, and 3.175 mm gauge hardware cloth walls. Each enclosure contained a 2 cm layer of chemical-free “play sand” for substrate (Quikrete 1113-51 Play Sand; ∼34 kg per enclosure), which lizards used for burrowing and thermoregulation. Enclosures also contained one basking site (a terracotta brick positioned lengthwise; 19.4 cm × 9.2 cm × 5.7 cm; length × width × height), and one shelter (20 × 20 cm wooden chipboard elevated 5 cm off the ground on wooden pegs). Between experimental batches, sand, basking sites, and shelters were replaced with new ones to reduce conspecific chemical signaling [21]. We installed aluminum flashing (7.6 cm × 40.6 cm) around the top perimeter of each enclosure to prevent the escape of crickets and lizards (Fig 1). Immediately after placing lizards into enclosures, we fed them one commercially sourced “medium” house cricket (*Acheta domesticus*) to standardize hunger levels at the start of a 48-hour acclimation period to the enclosure. During the acclimation period, researchers did not feed or interact with lizards except for a brief daily visit to check general welfare and habituation conditions in accordance with IACUC protocol.

### Experimental treatments

We designed experimental treatments to isolate the effects of blood sampling from potential background and handling-related effects. We administered a different treatment each day for three days, following a within-subject study design. We conducted a total of 54 trials, although only 52 were included in the final analyses because recordings from two trials were lost and dropped from the study (one from lizard ID 8 control treatment and one from lizard ID 9 immobilized treatment). To control for potential order and subject effects, treatments followed a Latin square design (Fig 2a). The order of individuals treated also followed a Latin square design to avoid treatment order effects (Fig 2b). All treatments were completed within 200 s (SEM = ± 8 s) of opening the enclosure. In the control treatment, we opened the enclosure, recorded dorsal body temperature, and remained in close proximity to the enclosure for the duration of the normal treatment period without otherwise interacting with the animal (Fig 1b). In the handling treatment, we lassoed the lizard, recorded ventral body temperature, and immobilized it (as if preparing it for a blood draw) for the duration of the treatment (Fig 1c). Finally, in the blood sample treatment, we lassoed the lizard and recorded ventral body temperature, then used a heparinized microcapillary tube (75mm length, inner diameter = 1.1 to 1.2mm) to collect 50 - 70uL of blood via the right retro-orbital sinus. After blood collection, we applied firm pressure on the sampled eye with a clean tissue to promote blood clotting. We observed no additional bleeding from lizards after treatment. We recorded treatment time from opening the enclosure to the return of the lizard onto the substrate.

**Fig 2.**
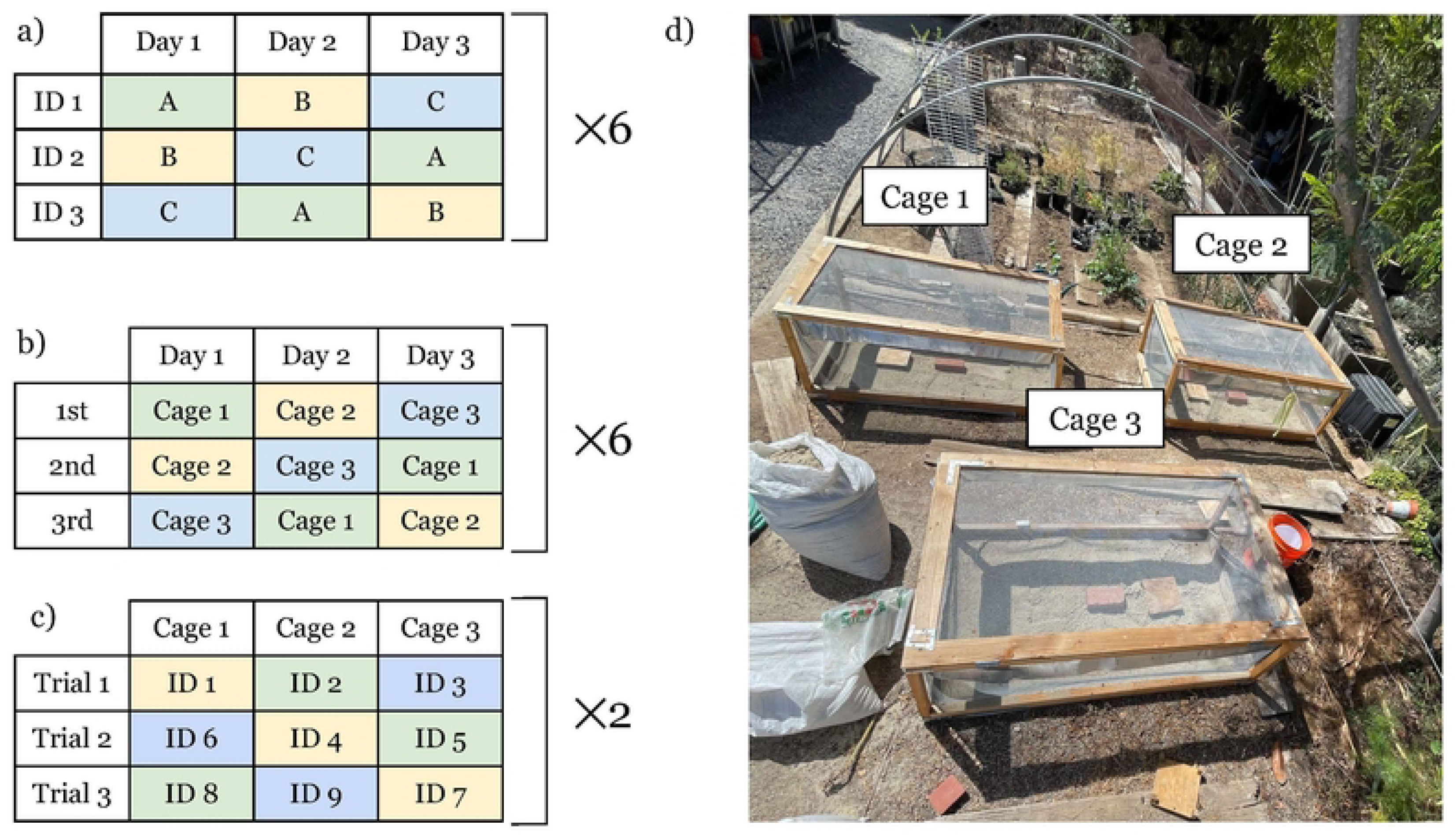
Experimental design and enclosure layout. a) Latin square design used to assign treatments to individuals (A = control, B = immobilization, C = blood sample) to account for possible treatment order effects; (b) Order of enclosure visits also followed a Latin square design to account for possible enclosure effects; (c) Enclosure assignment for each lizard (ID = lizard subject number); (d) Arrangement of enclosures in the facility. Enclosures were not moved during the experimental period.

To control for diel effects, we administered treatments between 1130 and 1530. To control for order and enclosure effects, enclosure assignments followed a Latin square design (Fig 2c). The first author (MJ) administered all treatments and served as the primary observer across all trials to eliminate observer bias.

### Feeding trials

Following each treatment, we returned the lizard to its enclosure and placed one cricket into the enclosure within 15 cm of the lizard’s head to initiate the feeding trial. We recorded three measures of feeding behavior: attack latency, feeding latency, and prey handling time. We defined attack latency as the time between prey introduction and the initiation of the first feeding attempt, which was characterized as a directed advancement towards the cricket (focused gaze and directional movement toward the prey, usually with mouth open). Feeding latency was the time from prey introduction to successful prey capture, characterized by the mouth closing around the cricket. In cases where lizards successfully captured prey on their first feeding attempt, attack latency was equal to feeding latency. We defined prey handling time as the time between prey capture to prey ingestion, which was marked by the cricket disappearing in the lizard’s mouth and the jaw remaining closed [22].

Feeding behavior was assessed for a maximum of 30 minutes post treatment. If five minutes elapsed with no lizard response to the cricket, which occurred when the cricket either moved out of the lizard’s visual range, hid in the shelter, buried, or crawled under flashing, a second cricket was added within 15 cm of the lizard’s head. In such cases, latency time was measured from the introduction of the first cricket regardless of which cricket was ultimately captured (Fig 1e) because the lizard had the opportunity to detect prey presence starting from the introduction of the first prey item. If 5 minutes elapsed after the addition of a second cricket with no response, no additional crickets were added and the assay ended at 30 minutes. We used Apple Voice Memo to take audio recordings of data from each lizard, and times were transcribed afterwards using a stopwatch. Upon the conclusion of each batch (3 lizards tested observed in each batch), lizards were marked on their dorsal side with non-toxic nail polish to prevent accidental recapture and released at the site of capture (Fig 1a) [23].

### Data analyses

We conducted all statistical analyses in R (v 4.4.0; [24]) and used mixed-effects Cox proportional hazards models (R package *coxme*) to test the effect of treatment on attack latency, feeding latency, and prey handling time. For attack latency, event status indicated whether a feeding attempt occurred (event status 0 = no feeding attempt, 1 = feeding attempt), for feeding latency whether prey was captured (0 = no prey capture, 1 = successful prey capture), and for prey handling whether prey was ingested (0 = not ingested, 1 = ingested). No lizards were excluded from the attack and feeding latency time-to-event analyses and Cox proportional hazards models. We visualized time-to-event using Kaplan-Meier curves (R package *survival*), with separate curves generated for each treatment within each feeding metric. Lizards that did not capture prey were excluded from prey handling analyses, as only lizards that successfully captured prey were capable of ingestion. When no feeding event occurred during the observation period, trials were assigned a time of 1800 s (the maximum assay duration) and treated as right-censored, meaning that the actual event was only known to be ≥ 1800 s. These right-censored values were included in time-to-event and Cox proportional hazards analyses but excluded from calculations of summary statistics. Observations were treated as independent events in the time-to-event analyses, with lizard ID included as a random effect in Cox proportional hazards models to account for the repeated sampling. Schoenfeld residuals indicated no violation of the assumption of proportional hazards for all models (p > 0.05). We ran additional Cox proportional hazards models including the order in which treatments were administered and enclosure assignments (see Fig 2d for enclosure arrangement) as covariates to assess potential confounding effects.

We then evaluated the persistence of treatment effects. We calculated the change in attack latency, feeding latency, and prey handling time between two consecutive treatments (24 h apart). Differences in metrics were calculated as the subsequent day minus the preceding day (e.g., attack latency on day 2 − attack latency on day 1. To further investigate whether the effects of blood sampling persisted into control treatments the following day, we performed Welch’s t tests to compare feeding metrics for lizards that received the control treatment first versus those that received the control treatment the day after blood sampling (control given day 2 or 3).

## Results

Across a 30-minute observation window, lizards had significantly longer attack and feeding latencies after blood sampling compared to immobilization and control treatments (Fig 3a & c). On average, attack latency measured 13.1 s (± 4.56) after the control treatment, 161.52 s (± 77.72) after the immobilization treatment, and 315.08 s (± 107.88) after blood sampling (Fig 3b; Table S2a). Lizards were 82.1% slower to attempt feeding after blood sampling compared to control conditions (hazard ratio (HR) = 0.179, p < 0.001) and 66% slower compared to the immobilization treatment (HR = 0.339, p < 0.01) (Fig 3a; Table S1a). Feeding latency followed a similar pattern: lizards averaged 13.13 (± 4.59) s after the control treatment, 162.86 (± 77.55) s after the immobilization treatment, and 542.64 (± 175.48) s after the blood sampling treatment (Fig 3d; Table S2b). Lizards were 88% slower to catch prey after blood sampling compared to the control treatment (HR = 0.112, p < 0.001) and 76% slower compared to immobilization (HR = 0.243, p < 0.01) (Fig 3c; Table S1b). Attack latency (HR = 0.529, p = 0.11) and feeding latency (HR = 0.46, p = 0.06) did not significantly differ between the immobilized and control treatments (Table S1a & b). Treatment order did not significantly affect attack latency, feeding latency, or prey handling time (all |z| ≤ 1.56, all p ≥ 0.12). Likewise, enclosure assignment had no significant effect on any feeding metric (all |z| ≤ 1.83, all p ≥ 0.07) (Table S5).

**Fig 3.**
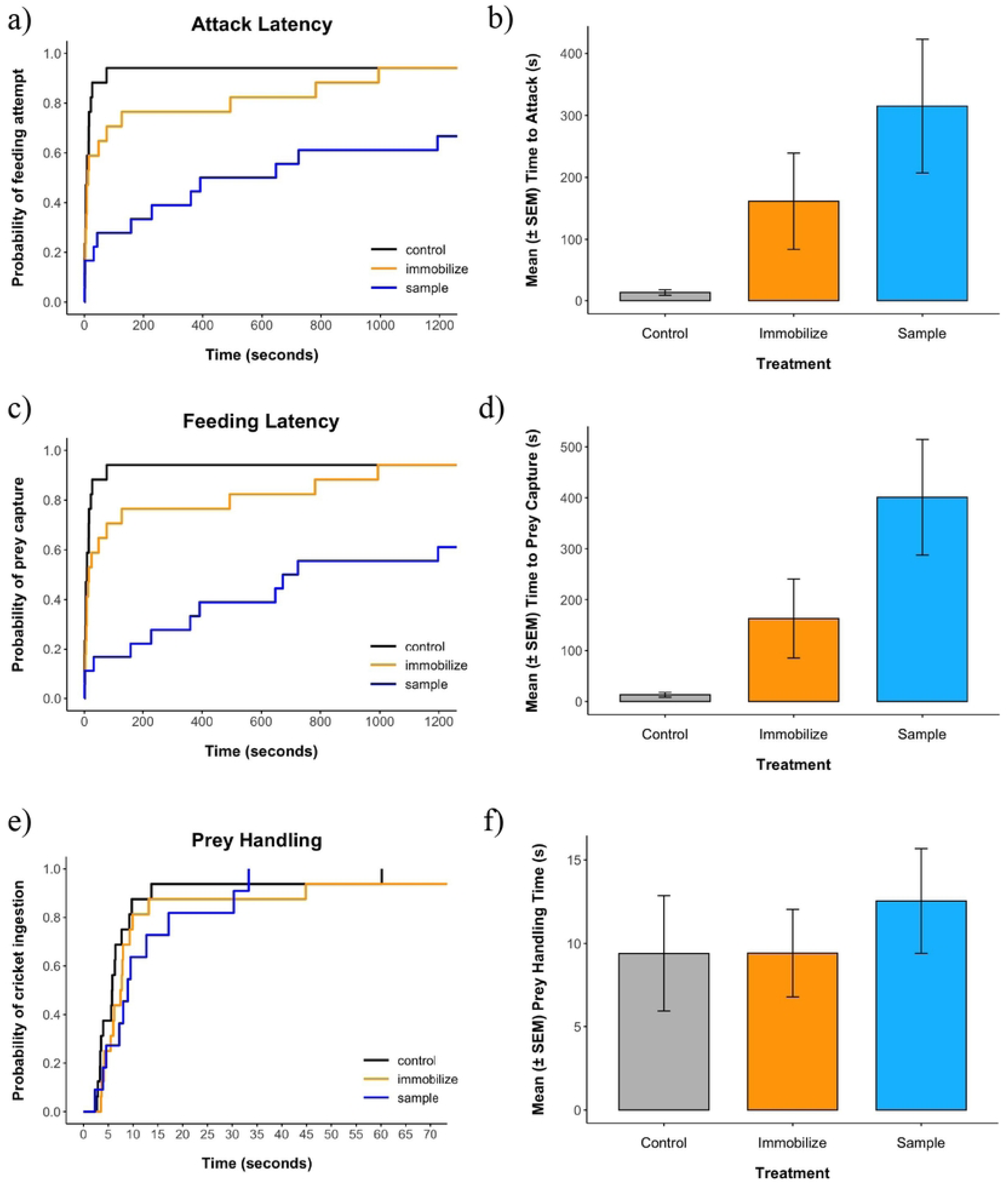
Time-to-event and mean feeding metrics in lizards subjected to control, immobilization, or blood sample treatment. (a-b) Lizards were slowest to make a feeding attempt after blood sampling (sampled versus immobilized HR = 0.42, p < 0.01; sampled versus control HR = 0.179, p < 0.001) and the mean attack latency was longest after blood sampling. (c-d) Rates to successfully capture prey (feeding latency) were also significantly slower after blood sampling compared to the immobilized (HR = 0.243, p < 0.01) and control (HR = 0.112, p < 0.001) groups. Kaplan-Meier curves for attack and feeding latency were truncated at 1200 s, as no feeding events occurred beyond these timepoints. (e-f) After successful prey capture, differences in prey handling rates were not significantly different between the three treatment groups (sampled versus immobilized HR = 0.876, p = 0.76; sampled versus control HR = 0.534, p = 0.16; immobilized versus control HR = 0.610, p = 0.20). Curves for prey handling time were truncated at 70 s for data visualization. One lizard (individual 5, immobilized treatment) ingested a cricket at 682.45 s and was excluded from the graph, but all observations were included in analyses. SEM = standard error of the mean.

Across all treatments, the rate at which lizards ingested prey after capture was not statistically different (blood sampled vs. immobilized HR = 0.876, p = 0.76; sampled versus control HR = 0.534, p = 0.16; immobilized versus control HR = 0.610, p = 0.20) (Table S1c). Mean prey handling time was 12.54 ± 3.14 s after blood sampling, 9.39 ± 3.46 s after the control treatment, and 9.42 ± 2.62 s after immobilization (Fig 3f; Table S2c).

We found no evidence that blood sampling effects persisted into the next day. Blood sampling, attack latency (median change = -136 s) and feeding latency (median change = -316 s) shortened under control conditions when measured the following day (Fig 4a & c, Table S3a & b). Prey handling time did not change in the 24 h between blood sampling and control treatments (median change = 0.26 s) (Fig 4e, Table S3c). Mean attack and feeding latency in day 1 control lizards did not differ from those in day 2 and day 3 control lizards (Welch’s t test: t = -0.36, df = 13.1, p = 0.73), nor was there a significant difference in mean prey handling time (t = -0.73, df = 10.17, p = 0.48) (Fig S4).

**Fig 4.**
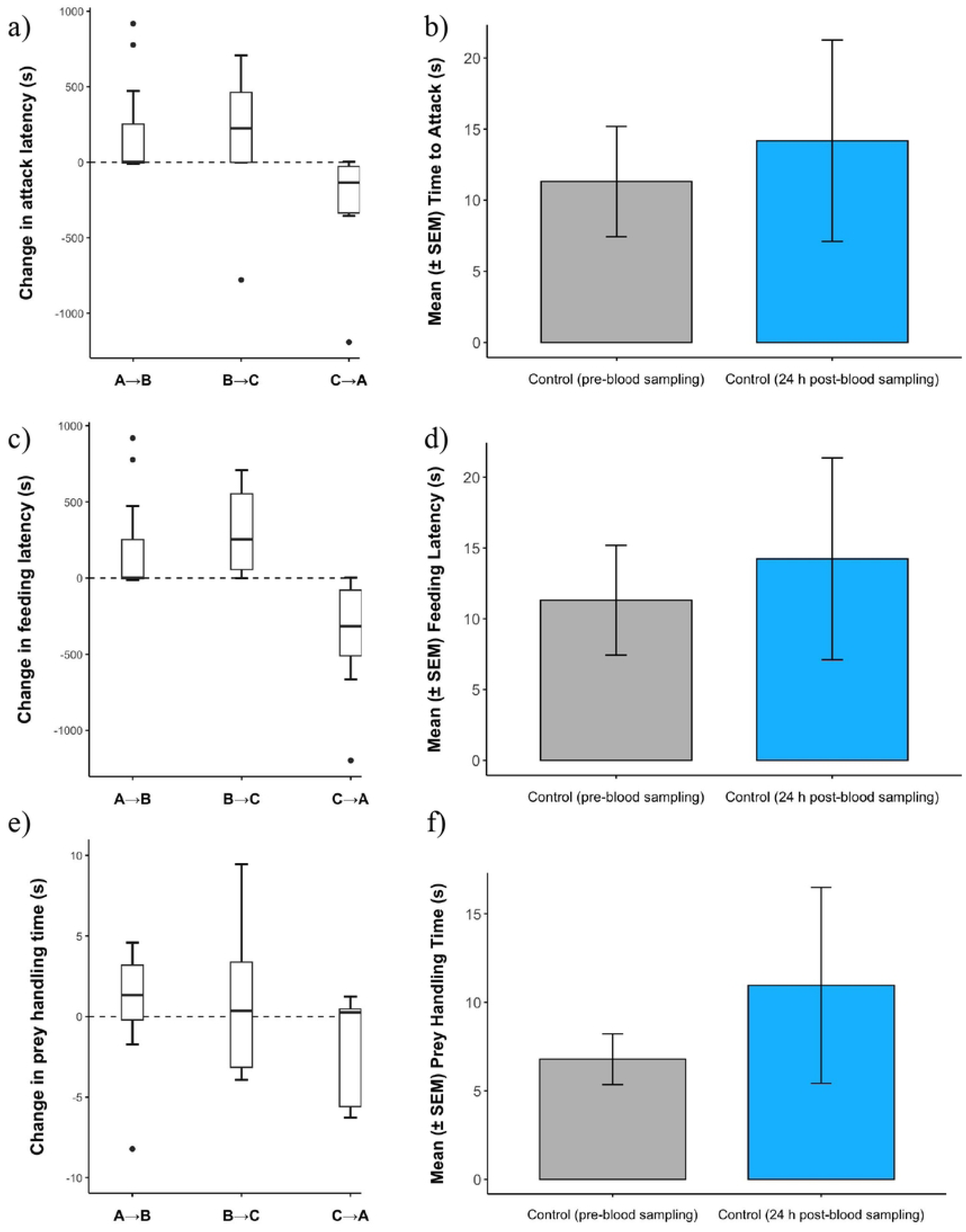
Persistence of affected feeding behavior across days (a, c, e), Differences in attack latency, feeding latency, and prey handling time between consecutive trial days (labelled on the x axes as “treatment given on previous day → treatment given on following day”) (A = control, B = immobilization, C = blood sample). Positive values indicated increased feeding metrics (longer attack latency, feeding latency, prey handling time) in the following treatment day, while negative values indicated decreased time measures. The median attack (n = 25) and feeding latency (n = 24) between control and immobilization treatment days (A → B) increased, and the same trend was demonstrated in the B → C lizards. Change in prey handling (n = 24) time did not exhibit a consistent trend. (b, d, f) The mean attack latency, feeding latency, and prey handling time in lizards where the control treatment was administered on trial day 1 (no treatments received beforehand) was slightly lower than those of lizards that were received the control treatment the day after blood sampling, but these differences between groups were not significant (attack latency p = 0.73, feeding latency p = -0.73, prey handling time p = 0.48).

## Discussion

Retro orbital blood collection remains a favored lizard blood sampling method for its convenience and presumed minimal invasiveness, serving as a foundational method in conservation and population genetics [6], disease vector ecology [1,2], systematics and evolutionary biology [25], and environmental endocrinology [3,5,7]. Despite its widespread use, current assumptions of the minimal welfare effects remain despite no formal test of its behavioral impacts ([26]; see [14] for physiological studies). Our study is the first to demonstrate a short-term reduction in feeding efficiency, which we define as the time it takes for prey to be captured and consumed, in western fence lizards after retro-orbital blood sampling. Our findings reveal important, previously neglected consequences of the method

Studies of lizard physiology permit the hypothesis that the lizard feeding reductions were indicative of stress. Physiological evaluations have documented elevated corticosterone levels, an established biomarker for stress in reptiles and vertebrates, following retro-orbital blood sampling [13,14] and cardiac puncture ([27,28]; but see [9,29] for exceptions). Although we did not measure corticosterone levels, reductions in feeding efficiency are a known symptom of stress in vertebrates, including reptiles [30,31]. If stress really was involved the behavior changes documented here, we may expect retro-orbital blood sampling to have additional physiological effects, such as to energetics, potentially leading to further welfare and fitness reductions.

In a similar vein, blood sampling may elicit an anti-predator response that entails reductions in feeding efficiency. Prolonged attack latency and decreased feeding rates have previously been identified as anti-predator behavior in broad-headed skinks [32] and alpine newts [33], and delayed feeding latency has been observed as neophobic behavior in geckos [34]. Aside from predator avoidance, decreased feeding rates confer no direct survival benefits and could be particularly detrimental to small lizards like *S. occidentalis* because their high metabolic rates per unit mass require efficient feeding to meet energy demands [33,35,36]. Sustained feeding reductions over the long term could lead to slower growth, reduced fertility, and overall impaired individual fitness [33]. However, negative effects on fitness may not necessarily arise if reductions in feeding are short term, as they were here [33,37].

An alternative hypothesis is that reduced feeding behavior resulting from retro-orbital blood collection is instead a result of impaired visual acuity. We observed two lizards (ID 8 and ID 14) rubbing their sampled (right) eye with either their forelimb, on the enclosure wall, or on a brick substrate immediately after a blood draw was taken, suggesting signs of eye irritation. However, no additional signs of visual impairment (e.g., effects on movement, higher number of failed prey capture attempts) were recorded in any other lizards.

Regardless of the specific basis of feeding reductions by fence lizards, its short-term nature suggests that retro-orbital blood sampling does not cause any prolonged harm or fitness impacts to lizards, at least in the context of a single blood sampling event. Consistent with our observations, alpine newts exhibited similar recovery from feeding reductions following handling and marking [38]. Return to normal feeding in reptiles is also indicated by documented habituation (e.g., alpine newts habituated to fish presence; [33]) or shifting to alternative prey or habitats [37], so there may be a general capacity among reptiles and amphibians to rapidly recover from acute stresses of human research procedures.

A major advantage of retro-orbital blood sampling over other blood sampling methods in reptiles is that it allows repeated blood collection over a short timeframe [5,39,40]. Such repeated sampling could dramatically alter the ability of animals to recover. Because we only assessed behavioral responses following a single retro-orbital blood draw, we caution against extending our conclusions to repeated sampling and encourage further investigation into the cumulative effects of serial retro-orbital blood draws. Allostasis theory posits that repeated responses to stressors can cause physiological dysregulation and deteriorate an organism’s ability to return to a baseline state, a phenomenon known as allostatic overload [41,42]. If serial blood draws compound reduced feeding behavior, and combined with prolonged activation of physiological stress responses, the method could contribute to allostatic overload, to the detriment of both welfare and survival [41,43].

Although immobilization of lizards did not alter feeding behavior relative to control conditions, handling effects were not completely absent. Lizards had a tendency for longer attack and feeding latencies after immobilization, but only feeding latency differences approached statistical significance. Although we did not detect significant handling effects, such effects have been documented in alpine newts [38] and other reptiles and vertebrates [12,14,44]. Therefore, the potential for handling effects should not be excluded from consideration.

Animal biologists continue to face longstanding challenges in considering how to utilize essential research methods while also minimally disturbing research subjects. Our findings of the short-term behavioral effects of a single retro-orbital blood draw indicate that this useful method can continue to be used as long as researchers take into account the impacts on feeding, and presumably other behaviors. However, the effects of repeated sampling remain unclear and need evaluation. Understanding the confounding effects of retro-orbital blood sampling on animal behavior can inform more judicious applications of the method, bolstering not only the welfare of animals we study but also the reliability of the insights we gain from them.

## Author Contributions

**May Jiang**: formal analysis (lead), funding acquisition (lead), data curation (lead), investigation (lead), writing (lead), review and editing (lead), conceptualization (equal), experimental design (equal). **David Daversa**: project administration (lead), funding acquisition (supporting), conceptualization (equal), experimental design (equal), formal analysis (supporting), investigation (supporting), review and editing (supporting). **H. Bradley Shaffer**: conceptualization (equal), experimental design (equal), review and editing (supporting), formal analysis (supporting).

## Acknowledgements

We would like to thank our collaborators in the Shaffer Lab and Blumstein lab for discussion and feedback on preliminary results; Max Reines for assisting in lizard and data collection; Ryan Page for constructing enclosures; Allison Keeney, Matthew Southall, Kyler Plouffe, and the staff at the UCLA Mathias Botanical Gardens for their general support; and the UCLA La Kretz Center, Stunt Ranch Reserve Research Awards Grant, and the Wild Animal Initiative for funding the work. We acknowledge the use of ChatGPT-4 in assistance with editing figures.

## Data Availability Statement

Original data and R scripts used for analyses are stored on Figshare at https://doi.org/10.6084/m9.figshare.33010886 and will be made publicly available at the time of manuscript acceptance.

